# Detecting diversifying selection for a trait from within and between-species genotypes and phenotypes

**DOI:** 10.1101/2023.10.02.559886

**Authors:** T. Latrille, M. Bastian, T. Gaboriau, N. Salamin

## Abstract

To quantify selection acting on a trait, methods have been developed using either within or between-species variation. However, methods using within-species variation do not integrate the changes at the macro-evolutionary scale. Conversely, current methods using between-species variation usually discard within-species variation, thus not accounting for processes at the micro-evolutionary scale. The main goal of this study is to define a neutrality index for a quantitative trait, by combining within- and between-species variation. This neutrality index integrates nucleotide polymorphism and divergence for normalizing trait variation. As such, it does not require estimation of population size nor of time of speciation for normalization. Our index can be used to seek deviation from the null model of neutral evolution, and test for diversifying selection. Applied to brain mass and body mass at the mammalian scale, we show that brain mass is under diversifying selection. Finally, we show that our test is not sensitive to the assumption that population sizes, mutation rates and generation time are constant across the phylogeny, and automatically adjust for it.

## Introduction

Determining whether a trait is under a particular regime of selection has been a long-standing goal in evolutionary biology. Fundamentally, distinguishing neutral evolution from selection requires determining which selective regime is supported by the observed variation of traits or sequences. The variation of phenotypes (traits) and genotypes (sequences) can be observed at different scales, across different development stages at the individual level, across different individuals and populations at the species level, and finally across different species at the phylogenetic level. All these systems require different assumptions and methodologies, and the endeavor to determine the selective regime for a given trait has thus incorporated theories, methods, and developments across various fields of evolutionary biology such as quantitative genetics, population genetics, phylogenetics and comparative genomics (Lynch & Walsh, 1998; Walsh & Lynch, 2018).

Leveraging individual variations within the same species, Genome-Wide Association Studies (GWAS) in humans have shown that traits are mostly polygenic (many loci associated with a given trait) and under stabilizing selection, while the loci affecting those traits are mostly pleiotropic (many traits associated with a given locus) with additive effects (Sella & Barton, 2019; Simons et al., 2018). Across several populations, by contrasting both trait and genetic differentiation, Q_ST_–F_ST_ methods have been used to determine the selective regime and to quantify the strength of selection acting on a trait (Leinonen et al., 2008; Merilä & Crnokrak, 2001). A trait differentiation (Q_ST_) higher than genetic differentiation (F_ST_) is interpreted as a signature of diversifying selection due to adaptation in different optimum trait value in the different populations (Lamy et al., 2012). Contrarily, Q_ST_ lower than F_ST_ is interpreted as a signature of stabilizing selection. However, Q_ST_–F_ST_ methods have been found to require many populations (O’Hara & Merilä, 2005), and that various factors can generate a spurious signal of selection (Edelaar et al., 2011; Pujol et al., 2008). Moreover, the test for diversifying selection is limited to recent local adaptation since the test is based on the variation observed within a single species. To disentangle selection from neutral evolution, trait variation can also be observed at a larger time scale. For example, change in mean trait value accumulates linearly with time of divergence from a sister species, and also proportionally to the trait variance (Lande, 1980a; Turelli, 1984). Empirically, this effect can be observed for genes with larger within-species variation in gene expression level, which exhibits a faster accumulation of divergence in mean expression level (Khaitovich et al., 2004). Altogether, both the trait variance and the evolution in mean value can be used to test for trait selection in a pair of species (Walsh & Lynch, 2018).

To disentangle neutral evolution and selection, trait evolution can also be observed at a larger time scale. For example, change in mean trait value accumulates linearly with time of divergence from a sister species, and also proportionally to the trait variance (Lande, 1980a; Turelli, 1984). Gene expression exhibits a similar accumulation as divergence in expression accumulates faster for genes with large within-species variation (Khaitovich et al., 2004). Altogether, both the trait variance and the evolution in mean value can be used to test for trait selection in a pair of species (Walsh & Lynch, 2018).

Alternatively, by accounting for the underlying relationships between several species, the selective regime for a quantitative trait can also be tested at the phylogenetic scale (Felsenstein, 1985). Under neutral evolution, the change in mean trait value along a given branch of the tree is normally distributed, with a variance proportional to divergence time (Hansen & Martins, 1996). As a result, the mean trait value can be modeled as a Brownian process branching at every node of the tree (Hansen & Martins, 1996; Harmon, 2018). Reconstructing the trait variation along the whole phylogeny as a Brownian process can thus constitute a null model of neutral trait evolution. Deviations from the assumptions of the Brownian process are however well known. When trait variation is constraint because of optimum mean trait values across or between species, the pattern of evolution can be modeled by the Ornstein-Uhlenbeck processes, which is often interpreted as a signature of stabilizing selection (Catalán et al., 2019). Alternatively, a trend in the Brownian process (the tendency of a trait to evolve in a certain direction without fixed optimum) is interpreted as a signature of directional selection at the phylogenetic scale (Silvestro et al., 2019). However, studies have shown that such comparative approaches are subject to different biases (Harmon, 2018). First, a trait under stabilizing selection for which the optimal trait value is also evolving as a Brownian process will not deviate from a Brownian process, and thus be wrongly classified as neutral (Hansen & Martins, 1996). In other words, the better fit of a Brownian process does not necessarily constitute proof of the neutral model. Second, even for a trait evolving under a neutral regime, the Ornstein-Uhlenbeck process might sometimes be statistically preferred over a Brownian process due to sampling artifacts (Cooper et al., 2016; Price et al., 2022; Silvestro et al., 2015). Those limitations, altogether with the use of mean trait estimates leaving out the variance in traits between individuals, easily generate misclassification of selection from methods at the phylogenetic scale.

At the frontier between micro and macroevolution, comparative methods at the phylogenetic scale have acknowledged the importance of modeling within-species variation together with changes in mean trait value to either describe measurement errors (Hansen & Bartoszek, 2012; Lynch, 1991), incorporate values for individuals (Felsenstein, 2008) or to scale the rate of change in mean trait value (Gaboriau et al., 2020; Gaboriau et al., 2023; Kostikova et al., 2016). Within-species variation has also been used to infer diversifying selection by estimating the ratio of between to within variation of many traits and test for deviation from the average ratio across traits (Rohlfs et al., 2014; Rohlfs & Nielsen, 2015). Here, our goal was again to use both variances between and within species to determine the selective regime of a quantitative trait. We build a novel framework that integrates trait variation at the phylogenetic and population scales together with estimates of molecular divergence at both scales. It allowed us to define an expected ratio of normalized variance between and within species while setting the threshold of this ratio for neutral, stabilizing, and diversifying selection. The ratio that we propose can be considered as a neutrality index for a any quantitative trait articulating trait and nucleotide variation within and between species. Importantly, our neutrality index also leverages nucleotide divergence and polymorphism to normalize trait variation at both scales, such that it does not require estimating population size (within-species) or speciation time (between species). From the field of population genetics, our study can be seen as the macro-evolutionary generalization of Q_ST_–F_ST_ methods to account for phylogenetic relationships between species. From the field of phylogenetics, our study can be seen as an alternative to the EVE model (Rohlfs et al., 2014; Rohlfs & Nielsen, 2015) for a single trait, where we set a threshold for neutral evolution by leveraging species nucleotide polymorphism and divergence.

## Materials and Methods

### Neutrality index for a quantitative trait

Prior to developing our neutrality index, we review theoretical expectations for variations of quantitative traits and genomic sequences under neutral evolution for both within- and between-species variation.

### Within-species trait variations

For a given trait, the genetic architecture is mainly defined by the number of loci encoding the trait (*L*) and the random additive effect of a mutation on the trait (*a*). New mutations are generating trait variance and the average effect of a mutation on the trait is 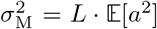. At the individual level, the mutational variance (*V*_M_) is the rate at which new mutations contribute to the trait variance per generation. As shown in Lande (1979, 1980b), *V*_M_ is a function of the mutation rate per loci per generation (*μ*) and 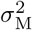 :

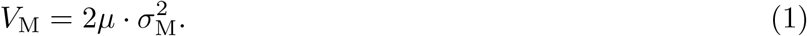

While in an infinitesimal model mutations supply new genetic variants, random genetic drift depletes standing variation (Barton et al., 2017; Sella & Barton, 2019; Turelli, 2017). For a neutral trait at equilibrium between mutation and drift (Lynch et al., 1998), the additive genetic variance in a species (*V*_A_) is a function of the mutational variance (*V*_M_) and the effective number of individuals in the population (*N*_e_):

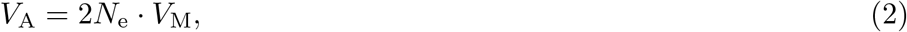

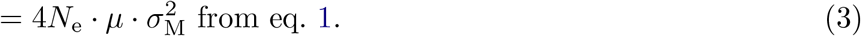

For any neutral genomic region of interest, the nucleotide diversity, *π*, is measured as the number of mutations segregating in the population divided by the length of the region. Any segregating mutations will eventually reach fixation or extinction due to random genetic drift and *π* is also at a balance between mutations and drift. As shown in Tajima (1989), *π* is a function of the mutation rate per loci per generation (*μ*) and the effective population size (*N*_e_):

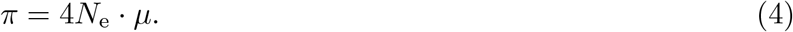

We define 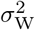 as the ratio of additive genetic variance of the trait (*V*_A_) over *π* of any neutral genomic region of interest. This ratio allows removing the effect of *N*_e_ and *μ*, which are parameters not related to the genetic architecture of the trait, giving 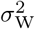 as a proxy of 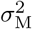 :

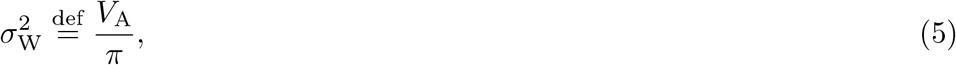

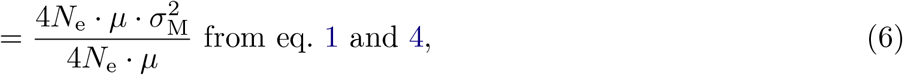

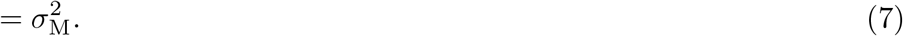

The additive genetic variance is also equal to the observed phenotypic variance (*V*_*P*_) multiplied by narrowsense heritability (*h*^2^; (Hill et al., 2008)), which leads to 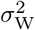 being a function of *V*_*P*_ and *h*^2^:

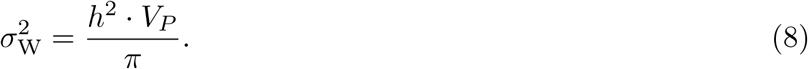

### Between-species trait variations

For a given species, we denote by 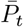 the mean value of the trait across the individuals in the species at generation *t*. If the trait is neutral and encoded by many loci as assumed by the infinitesimal model, 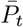 evolves as a Brownian process (Felsenstein, 1985; Hansen & Martins, 1996). The variance of 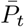 after *t* generations, Var 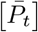 is given (Hansen & Martins, 1996) by:

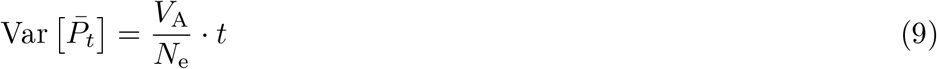

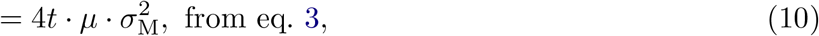

Moreover, for any genomic region under neutral evolution, some mutations will eventually reach fixation due to random genetic drift, resulting in a substitution of a nucleotide at the species level. The probability of fixation (ℙ_fix_) of a neutral mutation is 1*/*2*N*_e_ (Kimura, 1962). We can derive the substitution rate per generation *q* as the number of mutations per generation (2*N*_e_·*μ*) multiplied by the probability of fixation for each newly arisen mutations ℙ_fix_ (McCandlish & Stoltzfus, 2014), giving:

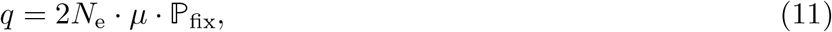

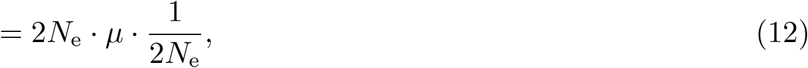

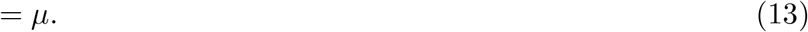

That is, if mutations are neutral, the rate of substitution within a genomic region equals the rate at which new mutations arise per generation for the same genomic region (Kimura, 1968).

After *t* generations and assuming that no multiple substitutions occurred at the same site, the nucleotide divergence *d*, which is the fraction of the genomic region that generated a substitution, will be *t* multiplied by the nucleotide substitution rate per generation (*q*):

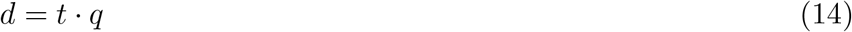

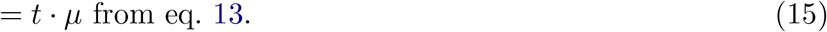

We define 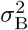 as the variance in the mean trait value (Var 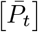) normalized by the nucleotide divergence of any neutral genomic region (*d*). This ratio allows removing the effect of *t* and *μ*, which are parameters not related to the genetic architecture of the trade, giving 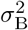 as another proxy of 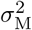 :

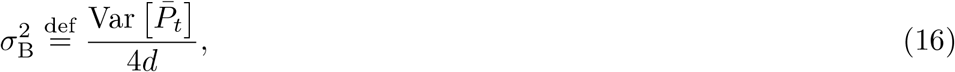

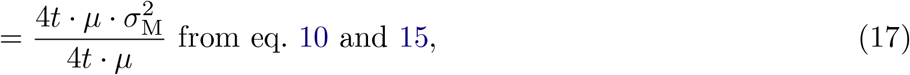

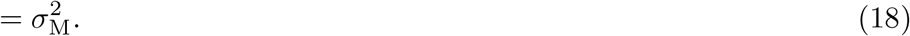

### Neutrality index

The variability between either individuals or species can be obtained for both quantitative traits and genomic sequences. At the population level, the variability of the trait between individuals can be combined with the nucleotide diversity of any neutrally evolving genomic region to obtain 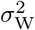, which equals 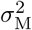 if the trait is neutrally evolving (see above). At the phylogenetic level, the variability of the mean trait value between species can be combined with the nucleotide divergence of any neutrally evolving genomic region to obtain 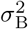 . Similarly, 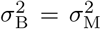 if the trait is neutrally evolving and the genetic architecture of the trait has not changed along the phylogenetic tree. We thus have, for a neutrally evolving trait:

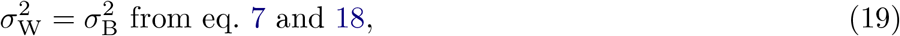

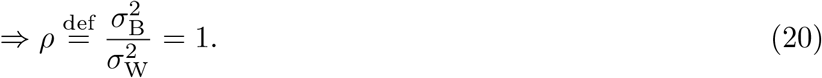

We define a neutrality index 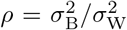 that will equal 1 for a trait evolving neutrally. Both 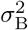 and 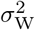 can be estimated using quantitative trait and genomic sequences within and between species, while neither the mutation rate (*μ*), nor the effective population size (*N*_e_) or time of divergence (*t*) need to be estimated. Moreover, the sequence from which *π* and *d* are estimated should be neutrally evolving, but they are not necessarily linked to the quantitative trait under study.

### Estimate

Based on the comparative framework that can account for phylogenetic inertia (Felsenstein, 1985; O’Meara et al., 2006), we provide a maximum likelihood estimate for *ρ* as well as a Bayesian estimate to derive posterior probabilities that the null model of neutrality (i.e. *ρ* = 1) is rejected.

### Maximum likelihood estimate

At the phylogenetic scale, for *n* taxa in the tree, ***D*** (*n × n*) is the distance matrix computed from the branch lengths (*d* as nucleotide divergence in units of substitutions per site) and the topology of the phylogenetic tree. The diagonal ***D***_*i,i*_ represents the total distances from the root of the tree to each taxon (*i*). The off-diagonal elements (***D***_*i,j*_ = ***D***_*j,i*_) are the distances between the root and the most recent common ancestor of taxa *i* and *j*. The state *P*_0_ at the root of the tree for the trait can be estimated from the *n ×* 1 vector of mean trait values 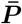 at the tips of the tree using maximum likelihood (O’Meara et al., 2006):

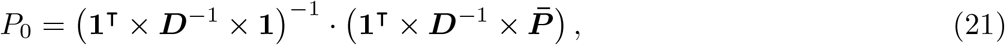

where **1** is an *n ×* 1 column vector of ones.

Finally, between-species variation 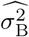 is estimated as (O’Meara et al., 2006):

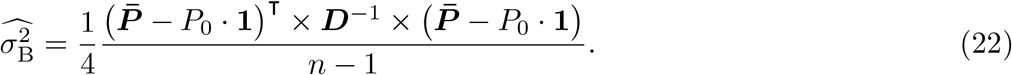

For a given species *i* with inter-individual data available, additive genetic variance of a trait (*V*_A,*i*_) is the product of heritability 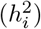 and phenotypic variance (*V*_*P,i*_). The ratio of *V*_A,*i*_ over nucleotide diversity of neutrally evolving sequences (*π*_*i*_) is a sample estimate of 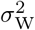 . Averaged across all species, we obtain the estimate 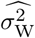 as:

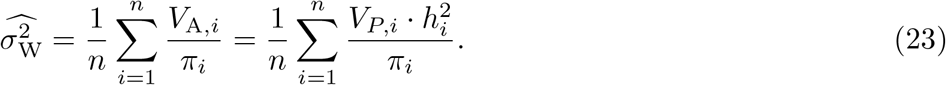

As depicted in fig. 1, the neutrality index is estimated as:

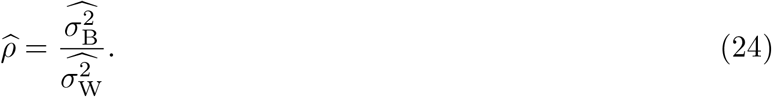

**Figure 1:**
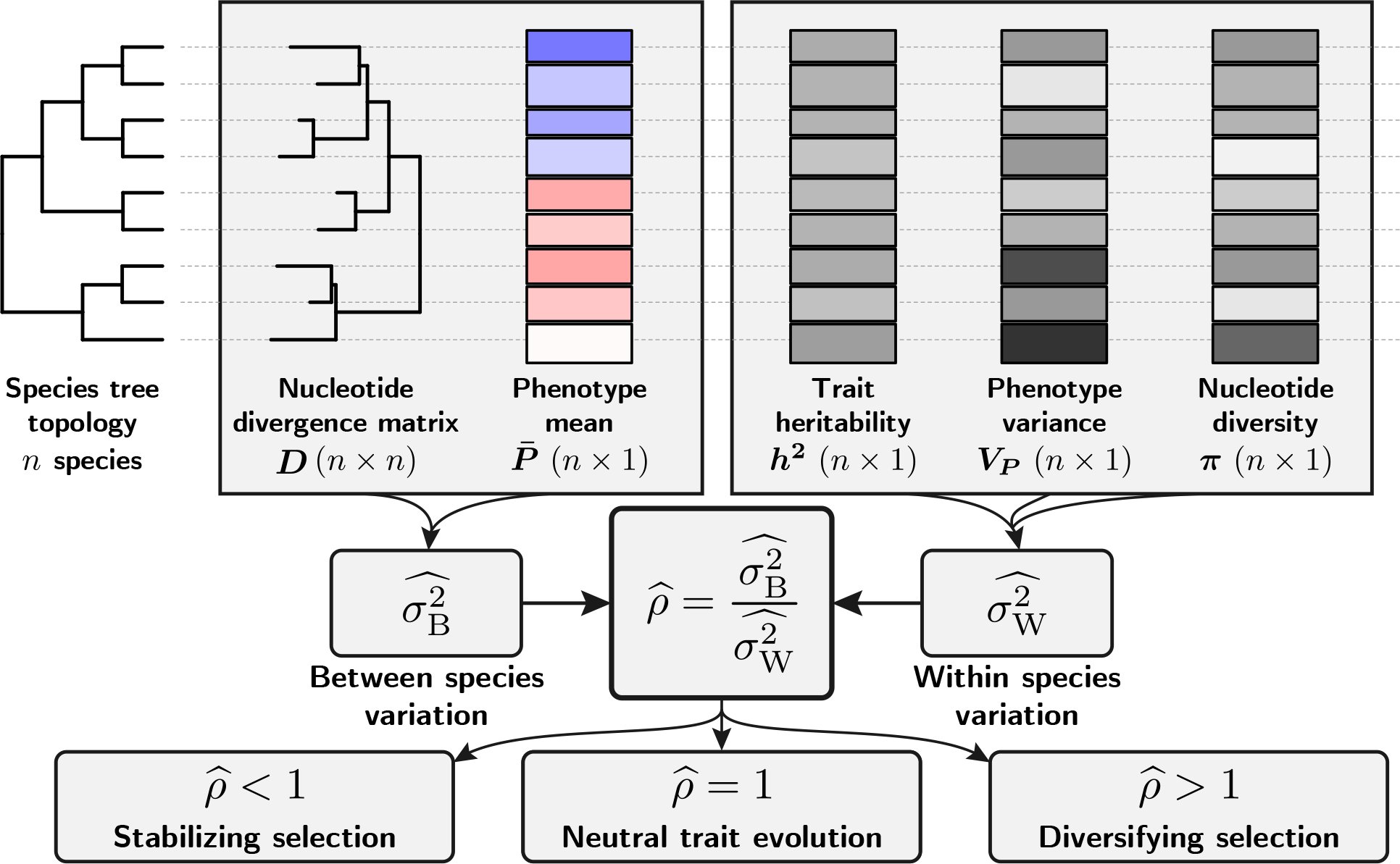
Between species, the change along the phylogeny of the mean phenotypic trait allows the estimation of between-species trait variation, 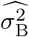, which is normalized by nucleotide divergence. Within species, the genetic variance allows the estimation of within-species trait variation, 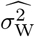, which is normalized by nucleotide diversity. 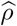 is the ratio of 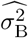 over 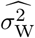 . Under neutral evolution, 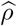 is expected to be equal to one. Under diversifying selection, the trait is heterogeneous between species, but homogeneous within species, leading to 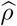 greater than one. Under stabilizing selection, the trait is homogeneous between species, leading to 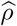 smaller than one. Importantly, the sequence from which nucleotide diversity and divergence are estimated should be neutrally evolving, but they are not necessarily linked to the quantitative trait under study, they allow for discarding the confounding effect on mutation rate diversity, population size and divergence time.

### Bayesian estimate

The Bayesian framework allows obtaining the posterior distribution of the neutrality index 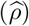 for a given trait. Even though 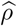 is estimated independently for each trait of interest in the maximum likelihood framework (previous section), here we generalize to *K* traits co-varying along the phylogenetic tree using the *BayesCode* software (Latrille et al., 2021). Trait variation along the phylogenetic tree is modeled as a *K*-dimensional Brownian process ***ℬ*** (1*×K*) starting at the root and branching along the tree topology (Huelsenbeck & Rannala, 2003; Lartillot & Poujol, 2011; Lartillot & Delsuc, 2012; Latrille et al., 2021). The rate of change of the Brownian process is determined by the positive semi-definite and symmetric covariance matrix between traits **Σ** (*K × K*). The off-diagonal elements of **Σ** are the covariance between traits, and the diagonal elements are the variance of each trait, thus corresponding to 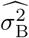 (see section S2.1). With an inverse Wishart distribution as the prior on the covariance matrix, the posterior on **Σ**, conditional on *ℬ* is also an invert Wishart distribution (see section S2.2). We used Metropolis-Hastings algorithm to sample ***ℬ***, while the posterior distribution of **Σ** is sampled using Gibbs sampling. For each trait and each species, the prior on heritability (*h*^2^) for each species is set as a uniform distribution with user-defined boundaries. Heritability and phenotypic variance for each trait are combined with nucleotide diversity to compute 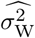 for each species before being averaged across species (as in eq. 23). From 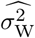 and **Σ**, the posterior distribution of 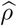 (as in eq. 24) is obtained for each trait. The posterior distribution of 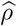 thus allows testing for deviation from neutrality (Fig. 1), for example, by computing 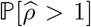 to test for evidence of diversifying selection and 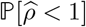 to test for evidence of stabilizing selection.

### Applicability to empirical data

Our method assumes that the narrow-sense heritability (*h*^2^) of a trait is known such as to estimate additive genetic variance (*V*_A_) from phenotypic variance (*V*_*P*_) as *V*_A_ = *h*^2^ · *V*_*P*_ . Fortunately, if heritability is not known, the test for diversifying selection can still be performed, although it is underpowered. Indeed, if the additive genetic variance is substituted by phenotypic variance, it is equivalent to assuming complete heritability (*h*^2^ = 1). Because *h*^2^ *≤* 1 by definition, we overestimate the within-species variation and thus underestimate 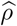. It is, however, possible to test for diversifying selection because testing for 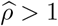 while using phenotypic variance instead of additive genetic variance means that knowing the additive genetic variance would have only increased the evidence for diversifying selection. Similarly, using the broad-sense heritability (*H*^2^) instead of narrow-sense heritability (*h*^2^) results in an underestimation of 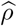 since *h*^2^ *≤ H*^2^. In contrast, the test for stabilizing selection is invalid if 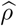 is underestimated. Several assumptions made by our test might not hold on empirical data and their consequences on the neutrality index and the test that can be performed are shown in Table 2.

### Simulation

We tested the performance of our neutrality index (*ρ*) to detect selection on a quantitative trait using simulations. We performed simulations under different selective regimes (neutral, stabilizing, diversifying), different demographic histories (constant or fluctuating population size) and different evolution of the mutation rate (constant or fluctuating). Simulations were individual-based and followed a Wright-Fisher model with mutation, selection and drift for a diploid population including speciation along a predefined ultrametric phylogenetic tree (Fig. 2A&B). Each individual phenotypic value was the sum of genotypic value and an environmental effect. The environmental effect was normally distributed with variance *V*_E_. We assumed that the genotypic value was encoded by *L* = 5, 000 loci, with each locus contributing an additive effect that was normally distributed with standard deviation *a* = 1 (Fig. 2A and section S1.1 for theoretical formulation). We assumed a trait with a narrow-sense heritability of *h*^2^ = 0.2 and computed the theoretical *V*_E_ accordingly (see section S1.1). Assuming a diploid panmictic population of size *N*_e_ = 50 at the root of the tree, and with non-overlapping generations, we simulated explicitly each generation along an ultrametric phylogenetic tree. For each offspring, the number of mutations was drawn from a Poisson distribution with mean 2 · *μ* · *L*, with the mutation rate per generation *μ*. From the empirical mammalian dataset (see next section), we computed an average nucleotide divergence from the root to leaves of 0.18 and average genetic diversity of 0.00276. We scaled parameters in our simulations to fit plausible values for mammals. We thus used a mutation rate of *μ* = 0.00276*/*4*N*_e_ = 1.38 *×* 10^*−*5^ per generation per locus and a total of *t* = 0.18*/*1.38 *×* 10^*−*5^ = 13, 500 generations from root to leaves, and the number of generations along each branch was proportional to the branch length.

**Figure 2:**
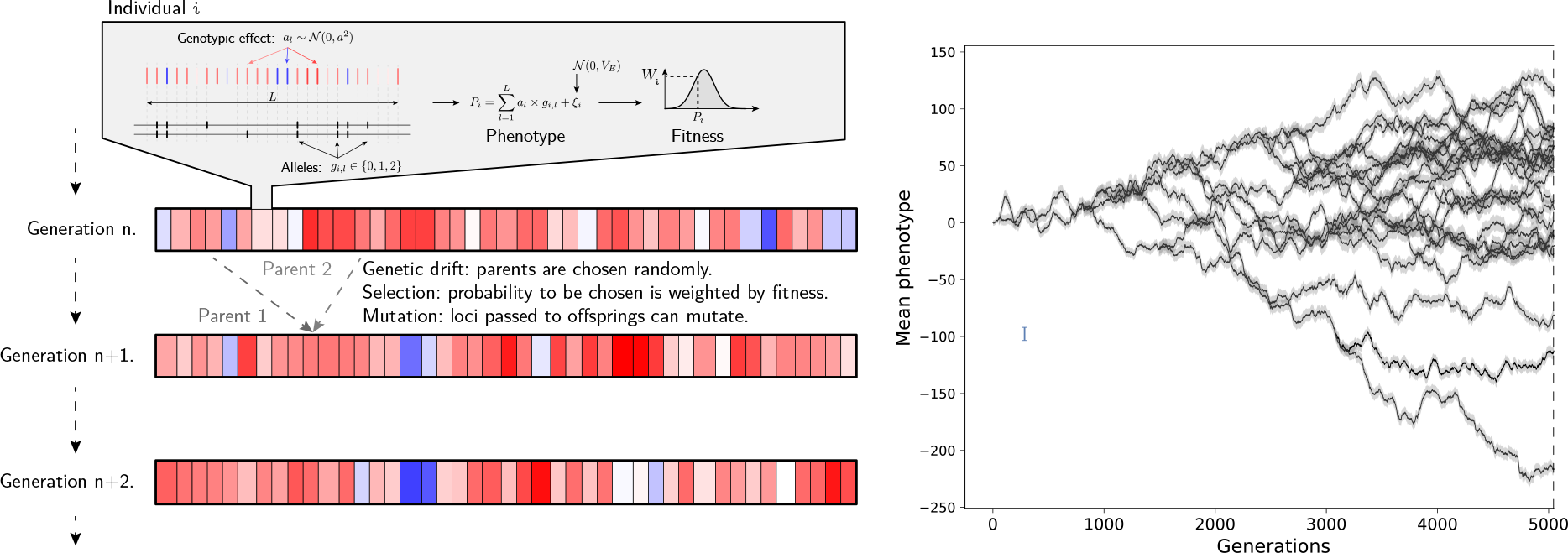
Wright-Fisher simulations with mutation, selection and drift. Left panel: For a given individual, the trait phenotypic value is the sum of genotypic value and a environmental effect (standard deviation *V*_E_). The trait’s genotypic value is encoded by *L* loci, with each locus contributing additively to the genotypic value. Parents are selected for reproduction to the next generation according to their phenotypic value, with a probability proportional to their fitness. Mutations are drawn from a Poisson distribution, with each locus having a probability *μ* to mutate. Drift is modeled by the resampling of parents. Right panel: examples of a trait evolving along a phylogenetic tree, with the mean phenotype (black line) and the variance of the trait genotypic value (gray area).

The changes in log-*μ* and log-*N*_e_ along the lineages were both modeled by a geometric Brownian process (*ℙ* (0, *σ*_*μ*_ = 0.0086) and 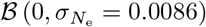, which led to a standard deviation of 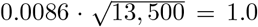 in log-space from root to leaves. An Ornstein-Uhlenbeck process was overlaid to the instant value of log-*N*_e_ provided by the geometric Brownian process to account for short-term changes between generations (OU 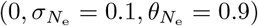). The geometric Brownian motion accounted for long-term fluctuations (low rate of changes 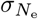 but unbounded), while the Ornstein-Uhlenbeck introduced short-term fluctuations (high rate of changes 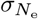 but bounded and mean-reverting). The simulation started from an initial sequence at equilibrium at the root of the tree and, at each node, the process was split until it finally reached the leaves of the tree. From a speciation process perspective, this was equivalent to an allopatric speciation over one generation.

A random genetic drift was introduced by resampling individuals at each generation, with each parent having a probability of being sampled that was proportional to its fitness (*W*). Selection was modeled as a onedimensional Fisher’s geometric landscape, with the fitness of an individual being a monotonously decreasing function of the distance between the individual and the optimal phenotype (Blanquart & Bataillon, 2016; Tenaillon, 2014). More specifically, the fitness of an individual was given by 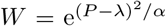, where *P* was the trait value of the individual, *λ* = 0.0 was the optimal trait value, and *α* = 0.02 was the strength of selection. Mutations were considered as a displacement of the phenotype in the multidimensional space. Beneficial mutations moved the phenotype closer to the optimum, while deleterious mutations moved it further away. Stabilizing selection was implemented by fixing the optimum phenotype to a single value (*λ* = 0.0). Diversifying selection was implemented by allowing the optimum phenotype to move along the phylogenetic tree as a geometric Brownian process (Hansen, 1997) (*λ ∼ℙ* (0, *σ*_*λ*_ = 1.0)). Neutral evolution was implemented by fixing the fitness landscape (*W* = 1), which meant that each individual had the same probability of being sampled at each generation.

Nucleotide diversity (*π*) was measured as the heterozygosity of neutral markers that were simulated along the phylogenetic tree but not linked to the trait simulated. Nucleotide divergence (*d*) was measured as the number of substitutions per site of neutral markers along the branches of the phylogenetic tree. The additive genetic variance was measured as phenotypic variance multiplied by heritability. Heritability was estimated from the slopes of the regression of offspring’s phenotypic trait values on parental phenotypic trait values (Lynch & Walsh, 1998) averaged over the last 10 simulated generations. Heritability was thus not a given parameter of the simulations, but rather measured as it would be in empirical data.

### Empirical dataset

We analyzed a dataset of body and brain masses from mammals. The log-transformed values of body and brain masses were taken from Tsuboi et al. (2018). We removed individuals not marked as adults and split the data into males and females due to sexual dimorphism in body and brain masses. We discarded species with only one representative sample. The mammalian nucleotide diversity was obtained from the Zoonomia project (Genereux et al., 2020), with nucleotide divergence obtained on a set of neutral markers in Foley et al. (2023), and with nucleotide diversity measured as heterozygosity in Wilder et al. (2023).

We also analyzed a dataset of primate species, with the nucleotide variation obtained from Kuderna et al. (2023) and the quantitative trait variation also from Tsuboi et al. (2018), using the same filtering as for the mammalian dataset. However, the primate nucleotide divergence was not obtained on a set of neutral markers as for the mammalian dataset, but across the whole genome.

## Results

### Neutrality index

For a neutral trait, the genetic architecture, meaning the number of loci encoding the trait and the average effect of a mutation on the trait, is formally related to both within and between-species variation of the trait. We defined the neutrality index as 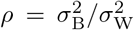, which equals 1 for a neutral trait (see Materials and Methods), suggesting that traits for which this relationship was not verified were putatively under selection. Under stabilizing selection, the variation between species is depleted because the mean trait value is maintained similar between different species, which leads to *ρ <* 1. In contrast, under diversifying selection, the variation between species is inflated because species will have potentially different trait values (Hansen, 1997), which leads to *ρ >* 1. Our neutrality index for a quantitative trait leveraged the data for any number of species, and took advantage of the signal over the whole phylogenetic tree, while at the same time taking into account phylogenetic inertia and addressing the non-independence between species (Fig. 1). This statistic was obtained as a maximum likelihood estimate 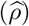, from eq. 23 and 22. We also devised a Bayesian estimate to obtain the posterior distribution of the neutrality index, and test for diversifying selection as 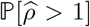, and stabilizing selection as 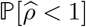.

Our neutrality index made a series of assumptions that we described in details in the Material and Methods section. Table 2 summarized these assumptions and outlined possible consequences for the neutrality test that we proposed.

### Results against simulations

The inference framework was first tested on independently simulated datasets matching an empirically relevant mammalian empirical regime (see Materials and Methods). Under constant population size (*N*_e_) and constant mutation rate (*μ*) across the phylogenetic tree (fig. 3, top row), we found no false negative for simulations of stabilizing (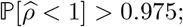 blue in fig. 3) or diversifying (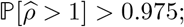 red in fig. 3) selection. For simulations under neutral evolution, 77% of those were correctly identified (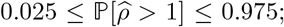 yellow in fig. 3), while 21% and 2% were wrongly detected as stabilizing or diversifying selection, respectively. Once we introduced fluctuating *N*_e_ and *μ* (Fig. 3, bottom row), our ability to identify simulations under either diversifying or stabilizing selection remained the same with all cases detected correctly. For simulations under neutral evolution, 51% of the simulations were correctly detected 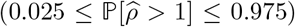, while 49% were detected as stabilizing selection 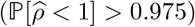 and none as diversifying selection.

**Figure 3:**
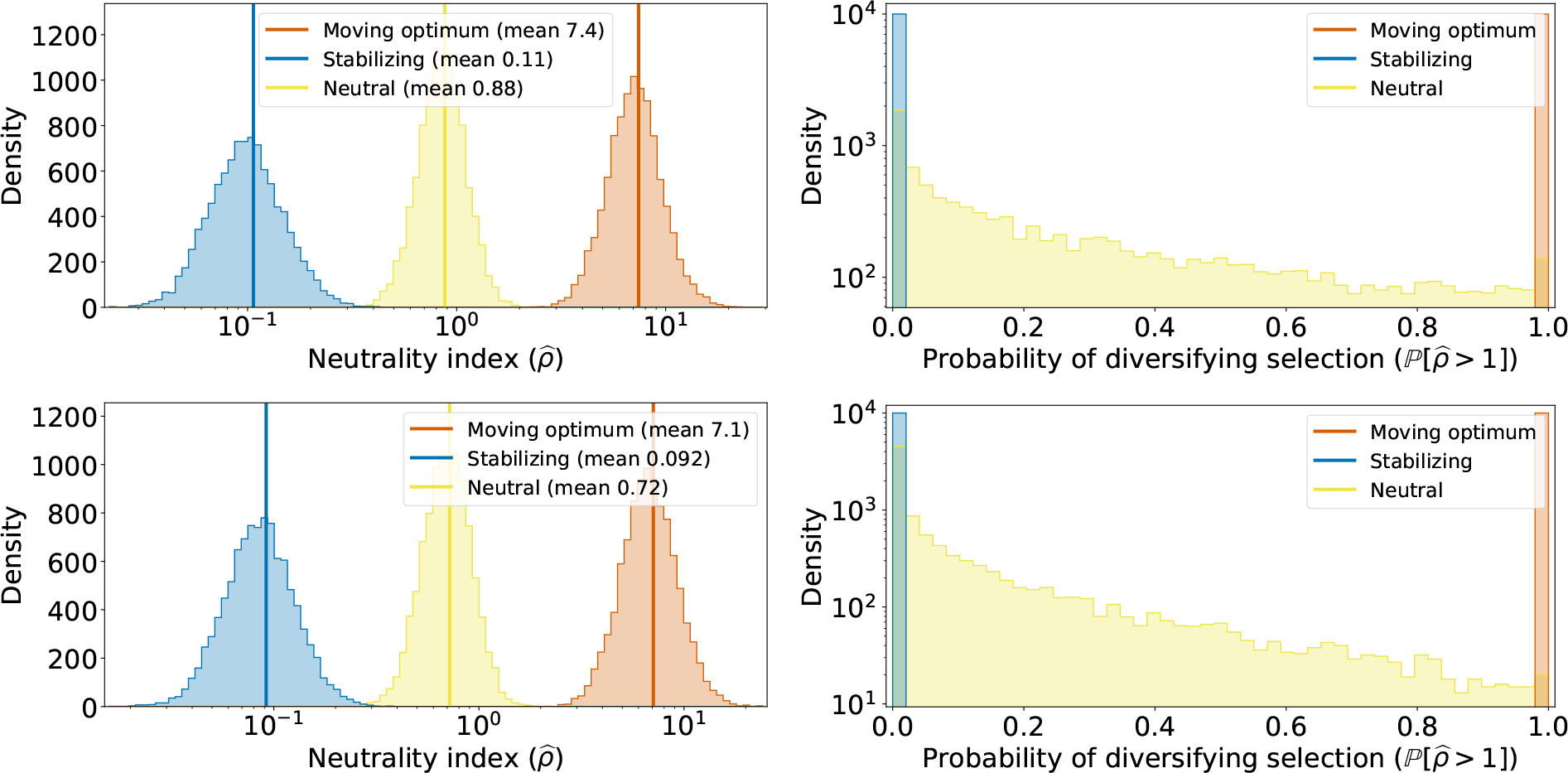
10, 000 simulations of trait evolution along a phylogenetic tree under different selection regimes. Traits simulated under stabilizing selection (blue), under a neutral evolution (yellow), and under a moving optimum (red). Histogram of ratio of between-species trait variation 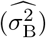 over within-species trait variation 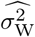 with 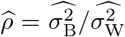 estimated from each simulated data (left) and probabilities of 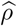 being greater than 1 (right). Effective population size (*N*_e_) and mutation rate (*μ*) were either constant (top row), or fluctuating as a Brownian process along the phylogenetic tree (bottom row).

### Results on empirical data

For mammalian body and brain mass, we obtained male (♂) and female (♀) trait variations. Combined with nucleotide diversity and divergence, we estimated 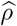 and posterior probabilities of diversifying selection under different assumptions for trait heritability as shown in the Table 1. Assuming complete heritability, brain mass was found to be under diversifying selection with posterior probabilities of 0.0 for both males and females. If we assumed that heritability (*h*^2^) of body mass was uniformly distributed between 20% and 40% (Hu et al., 2022), posterior probabilities of diversifying selection became 0.635 for males and 0.324 for females. Mammalian brain mass was found to be under diversifying selection with posterior probabilities of 0.877 for males and 0.972 for females when complete heritability was assumed. Assuming a uniform distribution between 20% and 40% for heritability led to posterior probabilities of diversifying selection of 1.0 for both males and females.

**Table 1:**
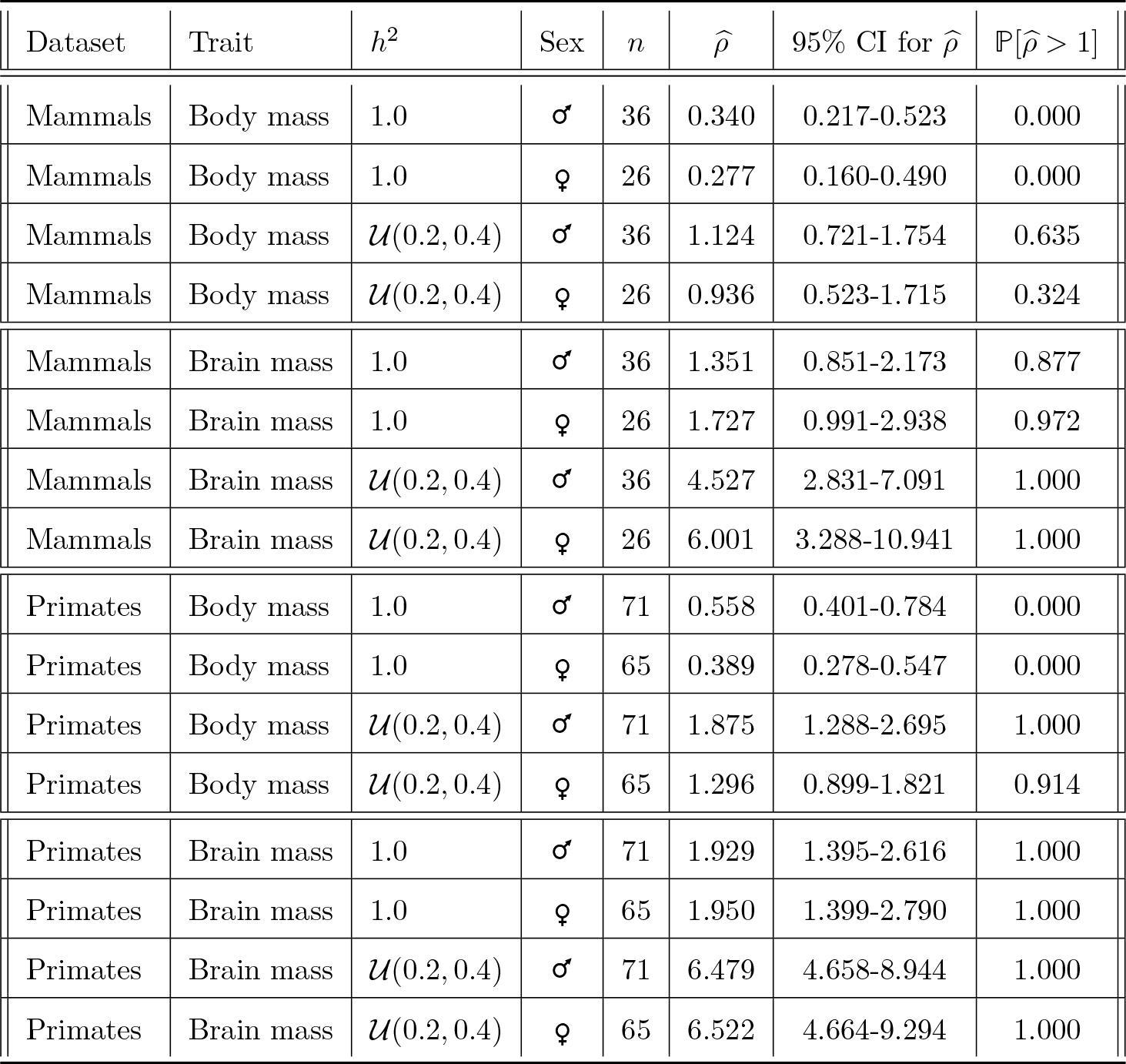
Test of diversifying selection on a mammal and a primate dataset, by splitting males (♂) and females (♀). Traits considered were body mass or brain mass (log-transformed). Heritability (*h*^2^) was either assumed complete (*h*^2^ = 1.0) or uniformly distributed between 20% and 40% (*h*^2^ *∼ U* (0.2, 0.4)). *n* was the number of species in the dataset. 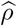 was the posterior estimate of our neutrality index, with the 95% credible interval (CI) for 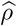 also computed. 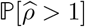 was the estimated posterior probability of diversifying selection.

| Dataset | Trait | $h^2$ | Sex | $n$ | $\hat{\rho}$ | 95% CI for $\hat{\rho}$ | $\mathbb{P}[\hat{\rho} > 1]$ |
| --- | --- | --- | --- | --- | --- | --- | --- |
| Mammals | Body mass | 1.0 | ♂ | 36 | 0.340 | 0.217-0.523 | 0.000 |
| Mammals | Body mass | 1.0 | ♀ | 26 | 0.277 | 0.160-0.490 | 0.000 |
| Mammals | Body mass | $\mathcal{U}(0.2, 0.4)$ | ♂ | 36 | 1.124 | 0.721-1.754 | 0.635 |
| Mammals | Body mass | $\mathcal{U}(0.2, 0.4)$ | ♀ | 26 | 0.936 | 0.523-1.715 | 0.324 |
| Mammals | Brain mass | 1.0 | ♂ | 36 | 1.351 | 0.851-2.173 | 0.877 |
| Mammals | Brain mass | 1.0 | ♀ | 26 | 1.727 | 0.991-2.938 | 0.972 |
| Mammals | Brain mass | $\mathcal{U}(0.2, 0.4)$ | ♂ | 36 | 4.527 | 2.831-7.091 | 1.000 |
| Mammals | Brain mass | $\mathcal{U}(0.2, 0.4)$ | ♀ | 26 | 6.001 | 3.288-10.941 | 1.000 |
| Primates | Body mass | 1.0 | ♂ | 71 | 0.558 | 0.401-0.784 | 0.000 |
| Primates | Body mass | 1.0 | ♀ | 65 | 0.389 | 0.278-0.547 | 0.000 |
| Primates | Body mass | $\mathcal{U}(0.2, 0.4)$ | ♂ | 71 | 1.875 | 1.288-2.695 | 1.000 |
| Primates | Body mass | $\mathcal{U}(0.2, 0.4)$ | ♀ | 65 | 1.296 | 0.899-1.821 | 0.914 |
| Primates | Brain mass | 1.0 | ♂ | 71 | 1.929 | 1.395-2.616 | 1.000 |
| Primates | Brain mass | 1.0 | ♀ | 65 | 1.950 | 1.399-2.790 | 1.000 |
| Primates | Brain mass | $\mathcal{U}(0.2, 0.4)$ | ♂ | 71 | 6.479 | 4.658-8.944 | 1.000 |
| Primates | Brain mass | $\mathcal{U}(0.2, 0.4)$ | ♀ | 65 | 6.522 | 4.664-9.294 | 1.000 |

**Table 2:** Assumptions breaks and their consequences on the estimation of within-species variation 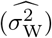, between-species variation 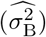, and on the neutrality index 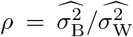 . The last two columns indicate whether the test for diversifying selection (*ρ >* 1) and for stabilizing selection *ρ <* 1 are conservative or invalid due to violated assumptions.

| Broken assumption | Consequences | $\widehat{\sigma}_W^2$ | $\widehat{\sigma}_B^2$ | Test $\rho > 1$ | Test $\rho < 1$ |
| --- | --- | --- | --- | --- | --- |
| Trait encoded by few loci | Between-species trait variation is underestimated | – | Underestimated | Conservative | Invalid |
| Sexual dimorphism | Within-species trait variation is overestimated | Overestimated | – | Conservative | Invalid |
| Inbreeding | Nucleotide diversity ($\pi$) is underestimated | Overestimated | – | Conservative | Invalid |
| Markers for polymorphism are negatively selected | Nucleotide diversity ($\pi$) is underestimated | Overestimated | – | Conservative | Invalid |
| Markers for polymorphism are positively selected | Nucleotide diversity ($\pi$) is underestimated | Overestimated | – | Conservative | Invalid |
| Markers for divergence are positively selected | Nucleotide divergence ($d$) is overestimated | – | Underestimated | Conservative | Invalid |
| Markers for polymorphism under balanced selection | Nucleotide diversity ($\pi$) is overestimated | Underestimated | – | Invalid | Conservative |
| Markers for divergence are negatively selected | Nucleotide divergence ($d$) is underestimated | – | Overestimated | Invalid | Conservative |
| Multiple nucleotide substitutions at the same locus | Nucleotide divergence ($d$) is underestimated | – | Overestimated | Invalid | Conservative |

We also analyzed a similar dataset for body mass focusing this time only at Primates (Table 1). For primates body mass, we found posterior probabilities of diversifying selection of 1.0 for males and 0.914 for females when assuming a uniform distribution for the heritability of body mass between 20% and 40%. Assuming complete heritability of body mass did not change the posterior probability for males, but increased the one for female to 1.0. Evidence for diversifying selection on body mass was therefore more pronounced in Primates than in mammals. However, the genetic markers used to normalize trait variance with nucleotide divergence were not necessarily neutral, which could create spurious false positives by artificially inflating 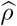 (Table 2 and methods).

## Discussion

In this study, we proposed a neutrality index for a quantitative trait that can be used within a statistical framework to test for selection. Our neutrality index for a trait, *ρ*, is calculated as the ratio of the normalized within-to between-species variation and it allowed the identification of the evolutionary regime of a quantitative trait. At the phylogenetic scale, trait variation between species was normalized by sequence divergence obtained from a neutral set of markers. Similarly, trait variation within species was normalized by sequence polymorphism obtained also from a neutral set of markers. Our estimate of 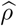 could be tested for deviation from the value of 1.0 expected under the null hypothesis of neutrality. Technically, the neutrality index can be estimated either as a maximum likelihood point estimate, or as a mean posterior estimate from a Bayesian implementation (see section S3). The latter also enabled the estimation of the posterior credible interval to test for departure from a neutrally evolving trait (e.g. 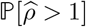). We tested our statistical procedure against simulated data and showed that our test was able to correctly detect simulations under diversifying selection (test of 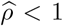) or under stabilizing selection (test of 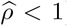). However, our test detected a spurious signal of stabilizing selection (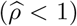) when we simulated the evolution of a neutral trait. We thus argue that our method should be used to detect diversifying selection, but that it had low accuracy to detect stabilizing selection due to false positives.

Our results showed that our method significantly improved over currently available methods to detect selection acting on a trait at the phylogenetic scale. Current methods relying on evolution of the mean trait value between species also tend to statistically prefer a model of stabilizing selection over a Brownian process when the trait is neutral (Cooper et al., 2016; Price et al., 2022; Silvestro et al., 2015). Our approach could in theory be applied to detect stabilizing selection at the phylogenetic scale, but we showed that it did not have the statistical power to identify those cases. In contrast, we showed that our method was able to identify correctly cases of diversifying selection, which is a clear an improvement over current methods that model only mean trait value. Indeed, under diversifying selection, mean trait value will not deviate from a Brownian process, and thus cannot be distinguished from neutral evolution (Hansen & Martins, 1996; Harmon, 2018). For example, testing the selective regime in the expression level of the majority of genes led to the selection of a Brownian process as the prefered model and the interpretation that the expression was evolving neutrally (Catalán et al., 2019). Our diversity index has the advantage to discriminate the alternative model of diversifying selection from the neutral case by comparing within- and between-species variation correctly normalized to remove confounding factors. Our approach is not the first one to normalize between-species variation to detect selection, but this was done by using within-species variations (Rohlfs et al., 2014; Rohlfs & Nielsen, 2015) and not estimates of neutral molecular divergence as done in our study. These studies have further compared their statistic across a pool of traits, which allowed them to identify outlier traits putatively under diversifying selection but without testing for selection on a single trait at a time (Gillard et al., 2021; Rohlfs & Nielsen, 2015). Instead, our procedure can be applied to a single trait, estimating the neutrality index and giving a statistical test for departures from the null model of neutral evolution for a single test. Our diversity index opens new avenues to revisit these studies and better test for the selective regime affecting the quantitative traits, assuming we have access to genomic datasets to estimate nucleotide divergence and polymorphism.

The main novelty of our study was to use the nucleotide divergence and polymorphism to normalize trait variation between and within species. In the context of within species variation, Q_ST_–F_ST_ tests have been developed to compare trait and sequence across several populations to test for selection (Leinonen et al., 2013; Martin et al., 2008). Our neutrality index also used the genetic sequences from which nucleotide divergence and polymorphism are estimated. Although the sequences should be neutrally evolving, they do not have to be necessarily linked to the quantitative trait under study. Nucleotide variation allows normalizing for diversity driven by confounding factors such as population sizes (*N*_e_), mutation rates (*μ*) and generation time (Hansen & Martins, 1996; Harmon, 2018). Thus our test avoids the estimation of the parameters, which are complex to correctly infer, and it also bypasses the estimation of divergence time, which was necessary in previous approaches (Walsh & Lynch, 2018). But importantly, by normalizing with sequence variation, we also showed using simulated data that our test was not sensitive to the assumption that *N*_e_, *μ* and generation time were constant across the phylogenetic tree, an unmet assumption empirically (Bergeron et al., 2023; Wilder et al., 2023). Indeed, under the neutral case of evolution, changes in *N*_e_, *μ* and generation time impacted similarly trait and sequence variation. The normalization by nucleotide divergence and polymorphism automatically absorbed long-term and short-term changes in *N*_e_, *μ* and generation time, which canceled out in the ratio of trait variation 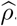.

Even though our test was developed for a quantitative trait, analogies with other tests of selection developed for molecular sequences also provided insight into its behavior. First, we acknowledge that our test took inspiration from the McDonald and Kreitman (1991) test devised for protein-coding DNA sequences, where synonymous mutations were used to determine the neutral expectation, and the inflation of divergence was compared to polymorphism within species. Second, because *ρ* was compared to 1, our test ultimately bear analogy to the codon-based test of selection, where the ratio of non-synonymous to synonymous substitutions (*ω*) is compared to 1 (Goldman & Yang, 1994; Muse & Gaut, 1994). As *ω <* 1 is interpreted as purifying selection acting on the protein, *ρ <* 1 is interpreted as stabilizing selection acting on the trait. Similarly, the interpretation of adaptation for *ω >* 1 is analogous to diversifying selection for *ρ >* 1. With this analogy in mind, we could leverage the vast literature discussing and interpreting the results of these tests and their pitfalls (Anisimova & Kosiol, 2009; Jensen et al., 2019; Nielsen, 2005). First, not rejecting the neutral null model of *ρ* = 1 did not necessarily imply that the trait was effectively neutral, since diversifying and stabilizing selection could compensate each other resulting in *ρ* = 1, analogously to *ω* = 1 under a mix of adaptation and purifying selection (Nielsen, 2005). Second, empirical evidence for *ρ <* 1 did not rule out diversifying selection, but rather that this diversifying selection was not strong enough to overcome the stabilizing selection, similarly to strong purifying selection resulting *ω <* 1 even though those genes and sites are under adaptation (Latrille et al., 2023). By explicitly modeling stabilizing selection as a moving optimum, it would theoretically be possible to tease apart the effect of diversifying and stabilizing selection in the context of quantitative traits to obtain a statistically more powerful test.

In the context of detecting diversifying selection on a trait, we argue that the main drawback of our method is that the additive genetic variance of the trait is required instead of the phenotypic variance. If phenotypic variance was used instead of additive genetic variance to estimate 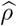, meaning that we assumed complete heritability, the neutrality index 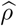 was ultimately underestimated. Similarly, using broad-sense heritability instead of narrow-sense heritability would result in underestimated 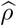. In such context, the test of stabilizing selection 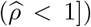 would be statistically invalid. However, the test of diversifying selection 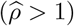 was underpowered although not invalided, meaning that absence of evidence would not be evidence of absence. As an example, even though we assumed complete heritability for brain mass, we uncovered diversifying selection in mammals since 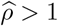.

The development of our neutrality index was also based on several assumptions that could be relaxed in future studies. First, we cannot predict the behavior of our test in the context of population structures, gene flow and introgression. These factors should be thoroughly investigated using simulations. Second, loci were assumed to contribute additively to the phenotype. Although the effects of dominance and epistasis is typically weak compared to the additive effects on the quantitative traits, their influence should be assessed (Crow, 2010; Hill et al., 2008). Third, the genetic architecture of the trait was assumed to be constant across the phylogenetic tree, whereas it might actually be variable among individuals and species (Huber et al., 2015; Tung et al., 2015). Such an assumption can theoretically be relaxed and changes in genetic architecture along the phylogenetic tree could jointly be estimated (Arnold et al., 2008; Gaboriau et al., 2020; Hohenlohe & Arnold, 2008; Kostikova et al., 2016). Finally, our Bayesian estimation could integrate uncertainty from the estimation of genetic variation, using sequences as input instead of estimated values of nucleotide diversity and divergence.

From an empirical point of view, our method required integrating genomic and trait variation, which could reduce the possible datasets to be used. However, such datasets will become more and more accessible and we showed the applicability of our method by applying it to the illustrative example of mammals brain and body mass. Because our test was also based on several assumptions that might not hold on empirical data, we also provided a table containing the main assumptions and their consequences on the neutrality index and the test that can be performed (Table 2). For example, at the primate scale, the evidence for 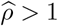 does not necessarily imply that the brain mass was evolving under diversifying selection since the markers used for nucleotide divergences were not neutral, which can lead to a spurious 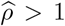. In conclusion, our study provided a statistical framework to test for diversifying selection acting on a quantitative trait while integrating the trove of genomic data available both within and between species, and we believe that our new approach is a promising tool to investigate the evolution of quantitative traits.

## Acknowledgements

We gratefully acknowledge the help of Nicolas Lartillot, Philippe Veber, Isabela Jeronimo do Ó, Anna Marcionetti, Julien Clavel, and Daniele Silvestro for their insightful discussions and Julien Joseph for his advice and reviews concerning this manuscript.

## Data and materials availability

The data that support the findings of this study are openly available in GitHub at github.com/ThibaultLatrille/MicMac. Snakemake pipeline, analysis scripts and documentation are available in the repository to replicate the study.

## Supplementary materials

### 1 Genetic architecture of the trait

#### 1.1 Genotype-phenotype map

- *L* is the number of loci encoding the trait.
- *a*_*l*_ *∼*𝒩 (0, *a*^2^) is the effect of a mutation on the trait at locus *l ∈ {*1, …, *L}*.
- *N*_e_ is the effective number of individuals.
- *g*_*i,l*_ *∈* {0, 1, 2} is the genotypic value at locus *l* for individual *i ∈* {1, …, *N*_e_}.
- 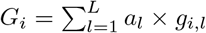 is the genotypic value for individual *i*.
- *ξ*_*i*_ *∼*𝒩 (0, *V*_E_) is the effect of environment on the trait for individual i.
- *P*_*i*_ = *G*_*i*_ + *ξ*_*i*_ is the phenotype for individual *i*.

**Figure S1:**
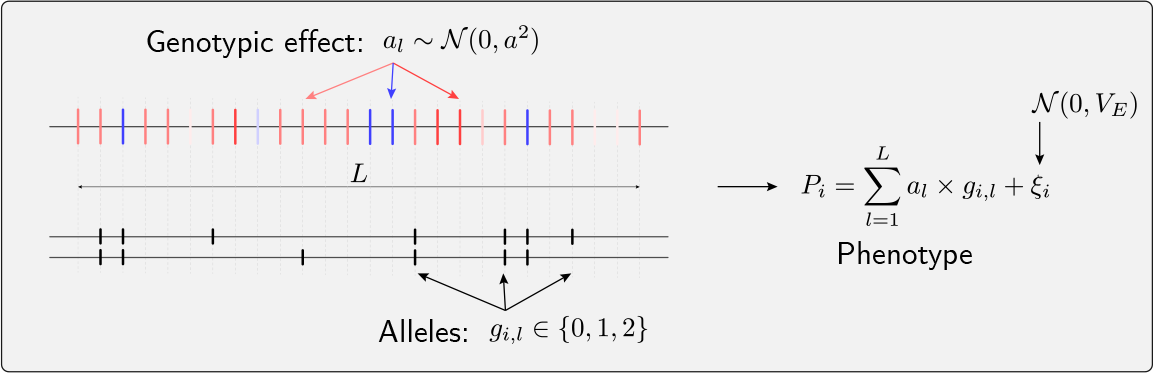
summary of trait’s genetic architecture.

within-species, the mean 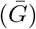 and variance (*V*_A_) of the genotype are:

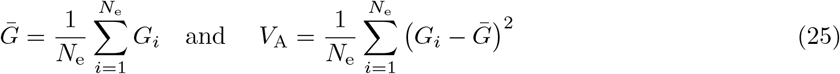

The theoretical additive genetic variance (*V*_A_) is a function of the number of loci (*L*) and the effect of a mutation (*a*) as:

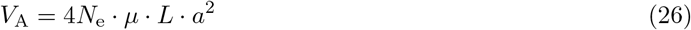

The mean 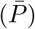 and variance (*V*_*P*_) of the phenotype are:

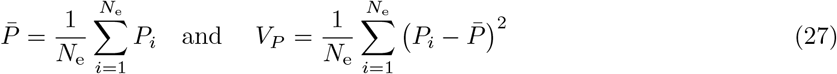

Heritability (*h*^2^) is defined as:

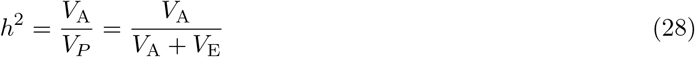

Altogether, effective population size (*N*_e_), the number of loci (*L*) and the effect of a mutation (*a*), we can compute the variance of the environment (*V*_E_) that is required to reach a given heritability (*h*^2^) as:

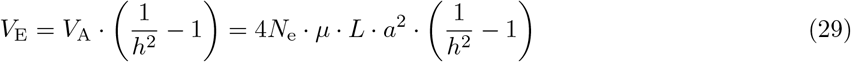

### 2 Bayesian estimate

#### 2.1 Multivariate Brownian process

Here we generalize to *K* traits evolving along the phylogeny and are correlated between them. Their variation along the phylogeny is modeled as a *K*-dimensional Brownian process ***ℬ*** (1 *× K*) starting at the root and branching along the tree topology. The rate of change of the Brownian process is determined by the positive semi-definite and symmetric covariance matrix between traits **Σ** (*K × K*). Along branch *j* with length *d*_*j*_, the Brownian process start at the ancestral node *𝒜* (*j*) with value ***ℬ*** (*𝒜* (*j*)), and ends at node *ℛ* (*j*) with value ***ℬ*** (*ℛ* (*j*)). The independent contrast ***C***_*j*_ defined as change in trait along the branch normalized by 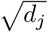 is a multivariate Gaussian:

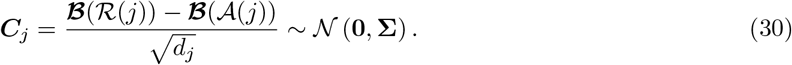

#### 2.2 Sampling the covariance matrix

From the independent contrast at each branch of the tree (***C***_*j*_), we can define the *K × K* scatter matrix, ***A***, as:

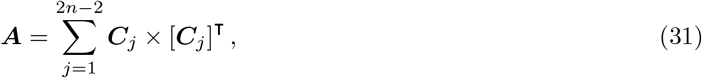

where 2*n −* 2 is the number of branches in the tree and *n* the number of taxa.

The prior on the covariance matrix is an inverse Wishart distribution, with *K* + 1 degrees of freedom:

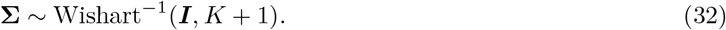

By Bayes theorem, the posterior on **Σ**, conditional on a particular realization of ***ℬ*** (and thus of ***C***) is an invert Wishart distribution, of parameter ***I*** + ***A*** and with 2*n* + 1 degrees of freedom.

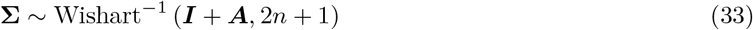

This invert Wishart distribution can be obtained by sampling 2*n* + 1 independent and identically distributed multivariate normal random variables ***Z***_*k*_ defined by

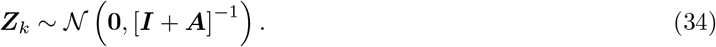

And from these multivariate samples, **Σ** is Gibbs sampled as:

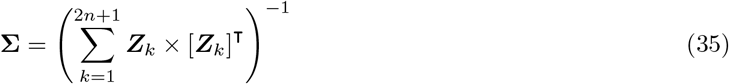

### 3 Bayesian and Maximum-likelihood implementation

Implementation is included within the *BayesCode* software, available at https://github.com/ThibaultLatrille/bayescode.

#### 3.1 Data formatting

Running the analysis on your dataset and compute posterior probabilities requires three files:

1. A phylogenetic tree in newick format, with branch lengths in number of substitutions per site (neutral markers).
2. A file containing the mean trait values for each species.
3. A file containing the variation within-species for each trait and the genetic variation within-species (neutral markers).

##### 3.1.1 Phylogenetic tree

The phylogenetic tree must be in newick format, with branch lengths in substitutions per site (neutral markers).

##### 3.1.2 Mean trait for each species

The file containing mean trait values for each species must be in a tab-delimited file with the following format:

| TaxonName | Body_mass | Brain_mass |
| --- | --- | --- |
| Panthera_tigris | 12.26 | 5.676 |
| Pithecia_pithecia | 7.256 | 3.436 |
| Colobus_angolensis | 9.176 | 4.284 |
| Saimiri_boliviensis | 6.845 | 3.279 |
| ⋮ | ⋮ | ⋮ |

The columns are:

- *TaxonName*: the name of the taxon matching the name in the alignment and the tree.
- As many columns as traits, without spaces or special characters in the trait.
- The values can be NaN to indicate that the trait is not available for that taxon.

##### 3.1.3 Trait variation for each species

The file containing trait variation for each species must be in a tab-delimited file with the following format:

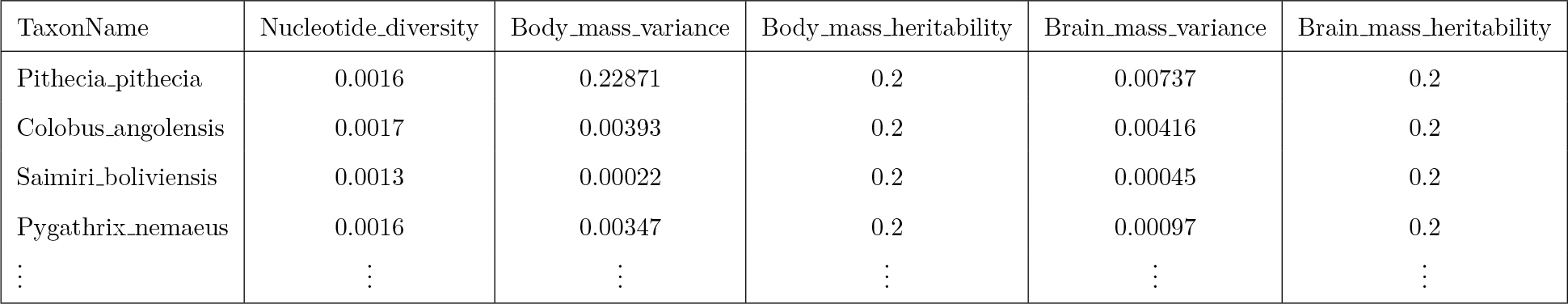

- *TaxonName*: the name of the taxon matching the name in the alignment and the tree.
- *Nucleotide diversity*: the nucleotide diversity within-species (neutral markers), cannot be NaN.
- As many columns as traits, without spaces or special characters in the trait.
- *TraitName variance*: the phenotypic variance of the trait within-species, can be NaN to indicate that the trait variance is not available for that taxon.
- *TraitName heritability* (optional): the heritability of the trait within-species, between 0 and 1, cannot be NaN.
- The columns with the `suffix variance and _heritability` are repeated for each trait.
- *TraitName heritability lower* (optional): the lower bound of the heritability of the trait within-species, between 0 and 1, cannot be NaN.
- *TraitName heritability upper* (optional): the upper bound of the heritability of the trait within-species, between 0 and 1, cannot be NaN.
- If the columns with the suffix `_heritability lower and heritability` upper are present, the heritability is randomly drawn from a uniform distribution between the lower and upper bounds.
- If the columns with the suffix `_heritability` is present, it is taken as is.
- If the additive genetic variance (instead of phenotypic variance) is available for a trait, the heritability can be omitted and will automatically be set to 1.0.

#### 3.2 Bayesian estimation

The executable nodetraits from *BayesCode* is used to run the Bayesian estimation of the model, and the executable readnodetraits is used to read the results.

Assuming that the file `data/body size/mammals.male.tsv` contains the mean trait values for each species, the file `data/body size/mammals.male.var` trait.tsv contains the variation within-species for each trait and the genetic variation within-species (neutral markers), and the file data/body size/mammals.male.tree contains the phylogenetic tree, the following commands are used to run the model and read the results.

##### 3.2.1 Running the model

`nodetraits` is run with the following command:

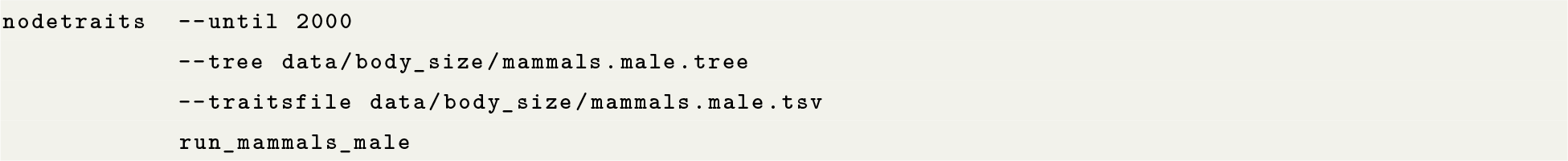

##### 3.2.2 Reading the results

Once the model has run, the chain run mammals male is used to compute the posterior distribution of the ratio of between-species variation over within-species variation with readnodetraits:

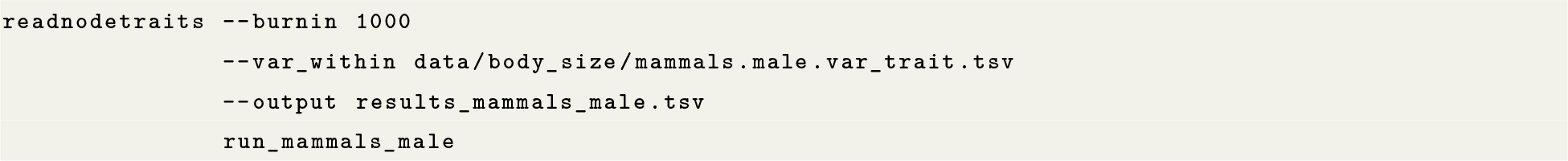

The file data empirical/chain name.ratio.tsv then contains the posterior mean of the ratio of between-species variation over within-species variation, the 95% and 99% credible interval, and the posterior probability that the ratio is greater than 1.

#### 3.3 Maximum likelihood estimation

To obtain the ratio (without the posterior credible interval and probability) using maximum likelihood computation, the following python script can be used:

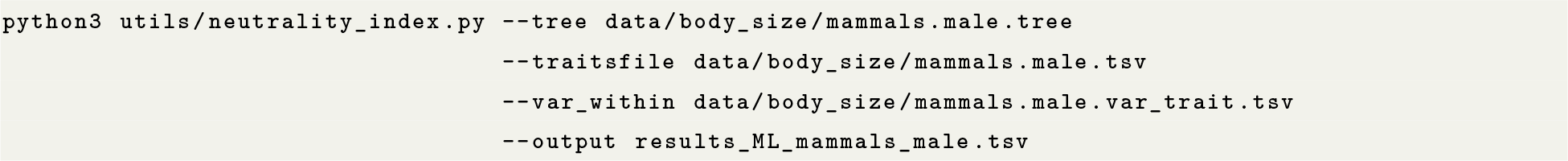

